# Nbs1 Mediates Assembly and Activity of the Mre11 complex

**DOI:** 10.1101/576082

**Authors:** Jun Hyun Kim, Alexander Penson, Barry S. Taylor, John H.J. Petrini

**Affiliations:** Molecular Biology Program, Memorial Sloan-Kettering Cancer Center, New York, NY10065, USA; Human Oncology and Pathogenesis Program, Memorial Sloan-Kettering Cancer Center, New York, NY10065, USA; Department Epidemiology and Biostatistics, Memorial Sloan-Kettering Cancer Center, New York, NY10065, USA; Center for Molecular Oncology, Memorial Sloan-Kettering Cancer Center, New York, NY10065, USA

## Abstract

We derived a mouse model in which a mutant form of Nbs1 (Nbs1^mid8^) exhibits severely impaired binding to the Mre11-Rad50 core of the Mre11 complex. The *Nbs1*^*mid8*^ allele was expressed exclusively in hematopoietic lineages (in *Nbs1*^*-/mid8vav*^ mice). Unlike *Nbs1*^*flox/floxvav*^ mice, which are Nbs1 deficient in the bone marrow, *Nbs1*^*-/mid8vav*^ mice were viable. *Nbs1*^*-/mid8vav*^ hematopoiesis was profoundly defective, exhibiting reduced cellularity of thymus and bone marrow, and stage specific blockage of B cell development. Within six months, *Nbs1*^*-/mid8*^ mice developed highly penetrant T cell leukemias. *Nbs1*^*-/mid8vav*^ leukemias recapitulated mutational features of human T-ALL, containing mutations in *Notch1, Trp53, Bcl6, Bcor*, and *Ikzf1*, suggesting that *Nbs1*^*mid8*^ mice may provide a venue to examine the relationship between the Mre11 complex and oncogene activation in the hematopoietic compartment. Genomic analysis of *Nbs1*^*-/mid8vav*^ malignancies showed focal amplification of 9qA2, causing overexpression of *MRE11* and *CHK1*. We propose that overexpression compensates for the meta-stable Mre11-Nbs1^mid8^ interaction, and that selection pressure for overexpression reflects the essential role of Nbs1 in promoting assembly and activity of the Mre11 complex.

## Introduction

The DNA damage response (DDR) is a network of pathways that mediate DNA repair and DNA damage signaling. The collective functions of the DDR preserve genomic integrity and viability in the face of myriad intrinsic and extrinsic genotoxic stressors. In addition, the DDR is a barrier to tumorigenesis (Ciccia and Elledge, 2010).

The Mre11 complex, composed of Mre11, Rad50, and Nbs1 is a sensor of DNA double-strand breaks (DSBs) and is required for the activation of the ATM axis of the DDR which parallels the ATR-Chk1 axis. The complex also plays an integral role in all aspects of DSB repair (Syed and Tainer, 2018).

Additionally, the Mre11 complex controls a non-canonical DNA damage response implicated in suppressing oncogene-driven carcinogenesis. Expression of the *neuT* oncogene in mouse mammary epithelium elicits the formation of DDR markers such as 53BP1 and γH2AX foci, as well as marked heterochromatic changes. In the context of Mre11 complex hypomorphism, those outcomes are not observed and the mice exhibit highly penetrant metastatic breast cancer. Deficiencies in other DDR components such as 53BP1 or mediators of apoptosis have little or no impact on the penetrance of *neuT*-induced malignancy in that experimental setting (Gupta et al., 2013).

Several lines of evidence also indicate that the Mre11 complex is critical for the process of DNA replication. Among them: Mre11 complex components are physically associated with the replication fork in normal as well as stressed conditions (Sirbu et al., 2013); the complex is required for viability of proliferating cells but dispensable for that of quiescent cells (Adelman et al., 2009); and the association of the complex with chromatin is qualitatively and quantitatively distinct in S phase compared to DNA damage induced association (Maser et al., 2001; Mirzoeva and Petrini, 2003). Collectively, these observations underscore the fact that with respect to preserving genomic integrity, DNA replication is an important nexus of regulation in normal as well as pathologic states, and that the Mre11 complex plays a central role at that nexus.

Whereas Mre11 and Rad50 orthologs are readily identifiable in all branches of life, Nbs1 (or Xrs2 in *S. cerevisiae*) orthologs are found only in Eukarya (Stracker and Petrini, 2011). Nbs1 interacts with Mre11 via a bipartite domain near its C terminus (Schiller et al., 2012). We previously derived a series of *Nbs1* alleles that targeted the Nbs1-Mre11 interface (called *Nbs1*^*mid*^ mutants for **M**re11 **i**nteraction **d**omain) so that the functionality of Nbs1 in isolation from the core Mre11-Rad50 complex could be assessed. The alleles that most severely impaired the interaction (*Nbs1*^*mid5*^ and *Nbs1*^*mid8*^) were lethal in MEFs and embryos. Conversely, a 108 amino acid fragment of Nbs1 that contained the bipartite Mre11 binding interface was sufficient to confer viability in MEFs as well as in the hematopoietic system (Kim et al., 2017). These data indicate that the essential function of Nbs1 is to stabilize and promote assembly of the active form of the Mre11-Rad50 core complex.

In this study, we used a hematopoietic cell specific *cre* (*cre*^*vav*^) (Stadtfeld and Graf, 2005) to test the hypothesis that the Nbs1^mid8^ gene product, which lacks four amino acids of the Mre11 interaction domain, specified sufficient residual function to support viability in the bone marrow. The hematopoietic compartment was chosen for this experiment because it is dispensable for embryogenesis and even severe deficits in hematopoietic development are compatible with postnatal viability in mice.

*cre*^*vav*^ was expressed in compound heterozygous mice containing floxed *NBS1* (*Nbs1*^*flox*^) (Demuth et al., 2004) and *Nbs1*^*mid8*^ alleles to establish mice expressing only *Nbs1*^*mid8*^ in hematopoietic lineages. In contrast to *Nbs1*^*flox/flox*^ mice, which upon *cre*^*vav*^ expression were Nbs1 deficient in the bone marrow and inviable, mice expressing *Nbs1*^*mid8*^ in hematopoietic lineages were viable postnatally, albeit with severe defects in hematopoiesis, and decrements in the levels of all hematopoietic components.

*Nbs1*^*mid8*^-expressing mice developed aggressive T cell malignancy which uniformly exhibited amplification and overexpression of *MRE11* as well as *CHK1* during the course of development and tumorigenesis. In *Nbs1*^*-/mid8*^ MEFs, co-overexpression of *MRE11* and *CHK1* was required for viability; overexpression of *MRE11* alone did not rescue *Nbs1*^*-/mid8*^ cellular lethality. The increased abundance of Mre11 appears to be required to mitigate the weakened interaction with Nbs1^mid8^ *in vivo*. These data support the interpretation that the essential role of Nbs1 to mediate assembly of the functional form of the Mre11 complex.

## Results

### *Nbs1*^*-/mid8vav*^ and hematopoiesis

Each member of the Mre11 complex is essential for viability at the cellular and embryonic levels. Genetic analyses of Mre11 and Rad50 have provided insights regarding their essential functions, whereas the essential role of Nbs1 has been less clear (Stracker and Petrini, 2011). Recently, we showed that the Mre11 interaction domain of Nbs1 is necessary and sufficient for cellular viability, and that Nbs1’s role is to promote assembly, activity, and nuclear localization of the Mre11 complex (Kim et al., 2017). In that study, a series of alleles in mice that impair the Nbs1 Mre11 interaction to varying extents were derived (*Nbs1*^*mid*^ mice).

The *Nbs1*^*mid8*^ allele, in which coding sequence for four amino acids is deleted from the Mre11 interaction domain (L**<u>KNFK</u>**KFKK; bold underlined amino acids deleted), abolishes ATM activation, was found to be embryonic lethal, and was unable to support the viability of immortalized murine embryonic fibroblasts (MEFs) (Kim et al., 2017). The Mre11 complex is required for hematopoiesis (Adelman et al., 2009; Balestrini et al., 2016; Callen et al., 2007; Reina-San-Martin et al., 2005). We asked whether the *Nbs1*^*mid8*^ gene product had sufficient residual function to allow hematopoietic stem cells expressing only *Nbs1*^*mid8*^ to support hematopoietic development. We chose hematopoiesis because although it is essential for postnatal viability, relatively severe defects are tolerated. And, we reasoned that amidst the abundant cellularity of the hematopoietic system, productive assemblies of Nbs1^mid8^ with the Mre11-Rad50 core might occur at a sufficient frequency to promote the development of at least a pauciclonal hematopoietic compartment if Nbs1^mid8^ retained some functionality.

*Nbs1*^*mid8*^ mice were crossed to *Nbs1*^*flox*^ mice (Demuth et al., 2004). The ensuing *Nbs1*^*flox/mid8*^ mice were then crossed to *vav*^*cre*^ in which *cre* expression is restricted to hematopoietic stem cells (Stadtfeld and Graf, 2005) to create *Nbs1*^*-/mid8*^ *vav*^*cre*^ mice (hereafter *Nbs1*^*-/mid8vav*^ mice). For the experiments described below, control mice include *Nbs1*^*flox/+*^ *vav*^*cre*^ (hereafter *Nbs1*^*-/+vav*^) and *Atm*^*floxlflox*^ *vav*^*cre*^ (hereafter *Atm*^*-/-vav*^). As expected, *Nbs1*^*flox/flox*^ *vav*^*cre*^ mice, in which hematopoietic cells are completely Nbs1 deficient exhibited perinatal lethality, succumbing at approximately two weeks to severe anemia (n = 13). In contrast, *Nbs1*^*-/mid8vav*^ mice were viable and born at normal Mendelian ratios, indicating that the *Nbs1*^*mid8*^ gene product is partially functional. *Nbs1*^*-/mid8*^ MEFs are inviable, and prior to death the cells exhibit severe genome instability and defective ATM activation (Kim et al., 2017). Hence, the viability of *Nbs1*^*-/mid8vav*^ bone marrow components was unexpected, and suggested the possibility of underlying compensatory genetic changes.

Hematopoiesis was severely impaired in *Nbs1*^*-/mid8vav*^ mice and did not phenocopy *Atm*^*-/-vav*^. Peripheral blood counts at six to eight weeks of age showed that both WBC and RBC numbers were decreased drastically in *Nbs1*^*-/mid8vav*^ compared to *Nbs1*^*-/+vav*^ mice, WBC; 8.66 ± 0.76 *vs.* 1.01 ± 0.14 (x10^6^/ml), RBC; 9.99 ± 0.38 *vs.* 2.51 ± 0.62 (x10^9^/ml) (Figure 1A and B). The percentage of both peripheral T cells and B cells also decreased in *Nbs1*^*-/mid8vav*^, T cell (%); 36.4, 26.7 *vs.* 0.4, 7.6, 9.8, B cell (%); 50.9, 58.4 *vs.* 2.9, 1.6, 8.3 (Figure 1D and E). *Nbs1*^*-/mid8vav*^ mice also exhibited a two fold elevation in platelet levels, likely a consequence of anemia (Figure 1C).

**Figure 1.**
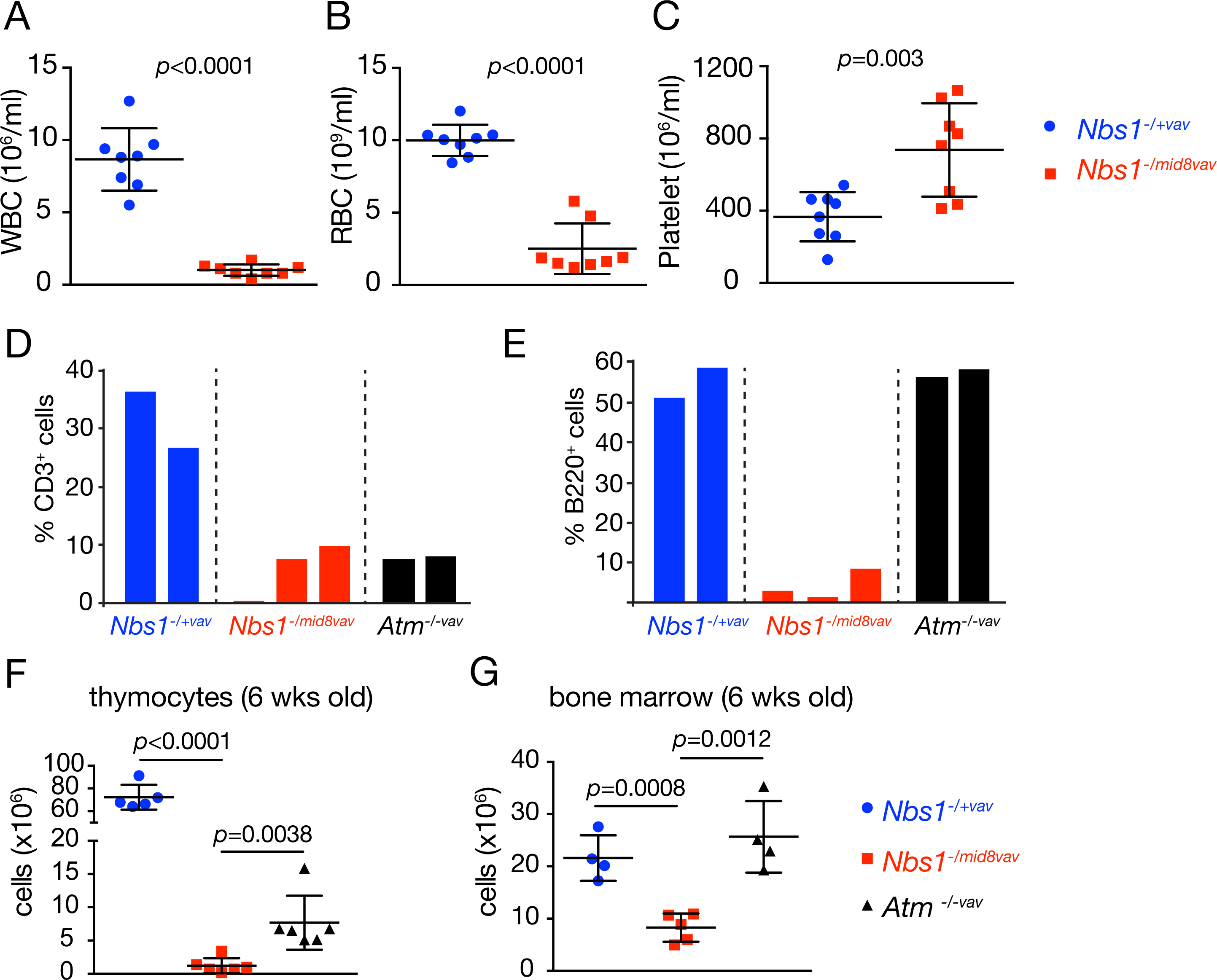
*Nbs11*^*-/mid8vav*^ allele leads to hematopoiesis failure. (A-C) Peripheral blood counts analysis of *Nbs1*^*-/mid8vav*^ mice at 6-8 week-old. (A) white blood cells (WBC), (B) red blood cells (RBC) and (C) platelet, *n=*8 mice of each genotype, mean ± SD, unpaired t test. (D) Percent CD3^+^ cells from peripheral blood. (E) Percent B220^+^ cells from peripheral blood. Each bar represents the data from individual mouse. (F) The number of thymocytes decreases in *Nbs1*^*-/mid8vav*^ mice, *n=*5 of *Nbs1*^*-/+vav*^, *n=*6 of *Nbs1*^*-/mid8vav*^, *n=*6 of *Atm*^*-/-vav*^, mean ± SD, unpaired t test. (G) The number of bone marrow cells decreases in *Nbs1*^*-/mid8vav*^ mice, *n=*4 of *Nbs1*^*-/+vav*^, *n=*5 of *Nbs1*^*-/mid8vav*^, *n=*4 of *Atm*^*-/-vav*^, mean ± SD, unpaired t test.

The cellularity of the thymus and bone marrow was also markedly decreased in *Nbs1*^*-/mid8vav*^ mice (Figure 1F and G). Analysis of *Nbs1*^*-/mid8vav*^ bone marrow was carried out at 6-8 weeks of age. The percentage of B lineage cells (B220^+^) and myeloid cell (Mac-1^+^ and Gr-1^+^ cells) were decreased in *Nbs1*^*-/mid8vav*^ mice compared to *Nbs1*^*-/+vav*^. B and myeloid lineages were reduced by roughly three fold (B: 25.03% ± 1.47 *vs.* 7.23 ± 0.92; myeloid 36.6% ± 2.83 *vs.* 14.32 ± 1.79). The levels of erythroid precursors (Ter119^+^ cells) were not changed (Figure 2A-C).

**Figure 2.**
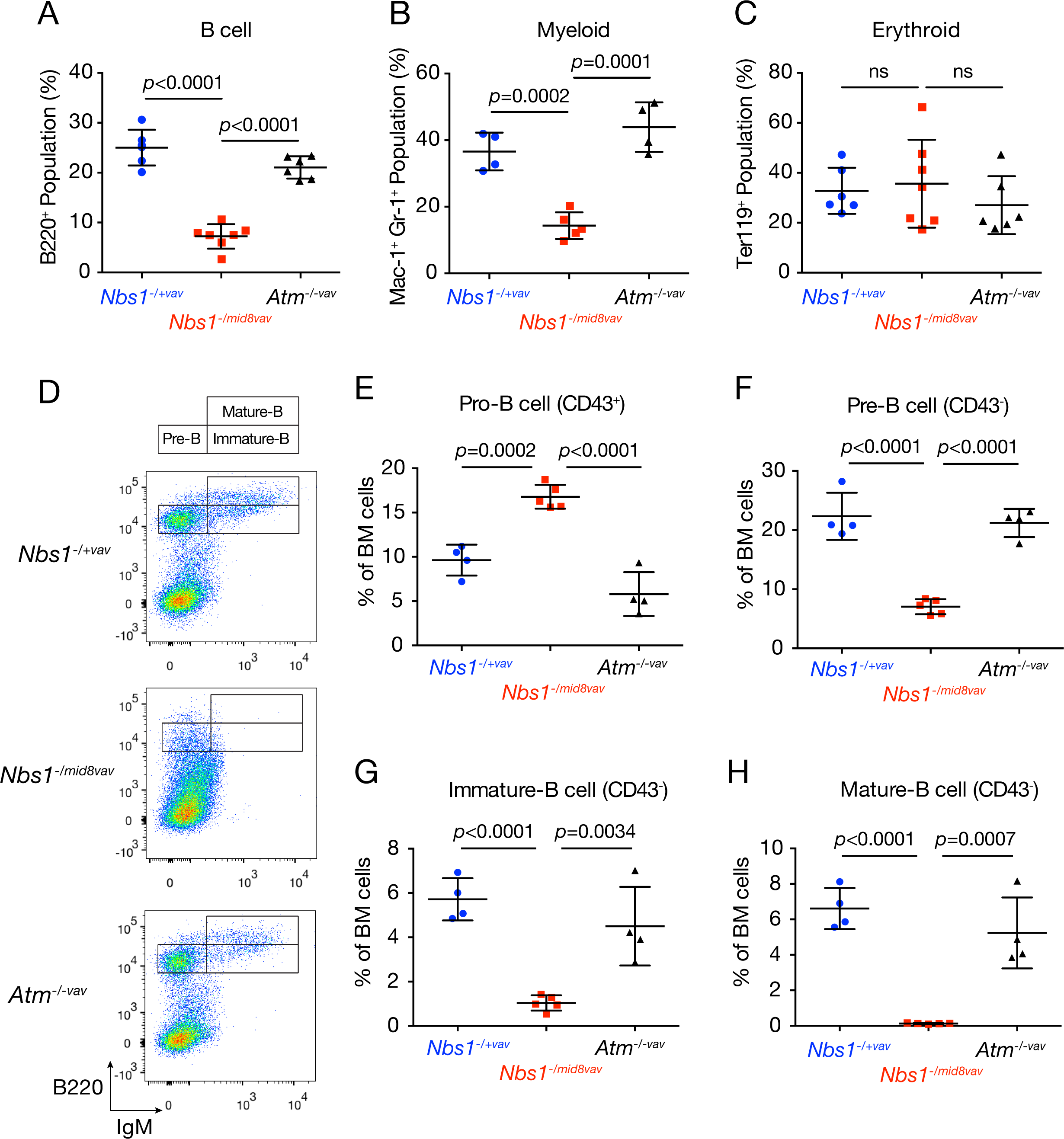
Depletion of B cell lineage cells in *Nbs1*^*-/mid8vav*^ bone marrow. (A) Percent B cell (B220^+^ cells) from bone marrow (BM) of indicated genotypes, *n=*5 of *Nbs1*^*-/+vav*^, *n=*7 of *Nbs1*^*-/mid8vav*^, *n=*6 of *Atm*^*-/-vav*^, mean ± SD, unpaired t test. (B) Percent myeloid cell (Mac-1^+^ and Gr-1^+^ cells) from BM of indicated genotypes, *n=*4 of *Nbs1*^*-/+vav*^, *n=*5 of *Nbs1*^*-/mid8vav*^, *n=*4 of *Atm*^*-/-vav*^, mean ± SD, unpaired t test. (C) Percent Erythroid cell (Ter119^+^ cells) from BM of indicated genotypes, *n=*6 of *Nbs1*^*-/+vav*^, *n=*7 of *Nbs1*^*-/mid8vav*^, *n=*6 of *Atm*^*-/-vav*^, mean ± SD, unpaired t test. (D) Representative scatter plot for B cell analysis of indicated genotypes. (E) Percent pro-B cells (CD43^+^) from BM. (F) Percent pre-B cells (CD43^-^, B220^+^, IgM^-^) from BM. (G) Percent immature-B cells (CD43^-^, B220^+^, IgM^+^) from BM. (H) Percent mature-B cells (CD43^-^, B220^+high^, IgM^+^) from BM cells. (E-H) Graphs show the data of indicated genotypes, *n=*4 of *Nbs1*^*-/+vav*^, *n=*5 of *Nbs1*^*-/mid8vav*^, *n=*4 of *Atm*^*-/-vav*^, mean ± SD, unpaired t test.

The depletion of B cell lineage cells appeared to coincide with onset of immunoglobulin gene assembly. Whereas pro-B cells (CD43^+^) from *Nbs1*^*-/mid8vav*^ were increased (Figure 2E), the levels of B220^+^ cells decreased beginning at the pre B stage when immunoglobulin heavy chain rearrangement commences (Alt et al., 2013; Teng and Schatz, 2015) to the immature B cell stage (CD43^-^), and IgM^+^ mature B cells were virtually undetectable (Figure 2F-H). These data suggest that *Nbs1*^*-/mid8vav*^ cells are unable to resolve DSBs formed during the course of B cell development, reminiscent of previous analyses of lymphocyte development in Mre11 complex mutants (Balestrini et al., 2016; Callen et al., 2007; Deriano et al., 2009; Helmink et al., 2009). The hematopoietic phenotype of *Nbs1*^*-/mid8vav*^ mice is distinct from that of *Atm*^*-/-vav*^ in all respects (Figure 1E-G, and Figures 2A-H), underscoring the point that the *Nbs1*^*-/mid8vav*^ phenotype reflects both the loss of both Mre11 complex DSB repair functions and ATM activation.

### *Nbs1*^*-/mid8vav*^ mice develop T cell lymphoma

In light of the predisposition to thymic lymphomas in *Atm*^*-/-*^ mice, we monitored a cohort of *Nbs1*^*-/mid8vav*^ mice for 12 months to assess the risk of malignancy. With 5.6 month of mean tumor free survival, 90% (18/20) of *Nbs1*^*-/mid8vav*^ developed hematologic malignancy, a significantly higher penetrance than that of *Atm*^*-/-vav*^ (Balestrini et al., 2016) (Figure 3A). In addition to exhibiting higher penetrance, the malignancies were distinct from those arising in *Atm*^*-/-vav*^. Most *Nbs1*^*-/mid8vav*^ tumors (17/18) were aggressive T cell lymphomas or leukemias (Table 1) which became disseminated to the spleen (Figure 3B ii and iii).

**Table 1.**
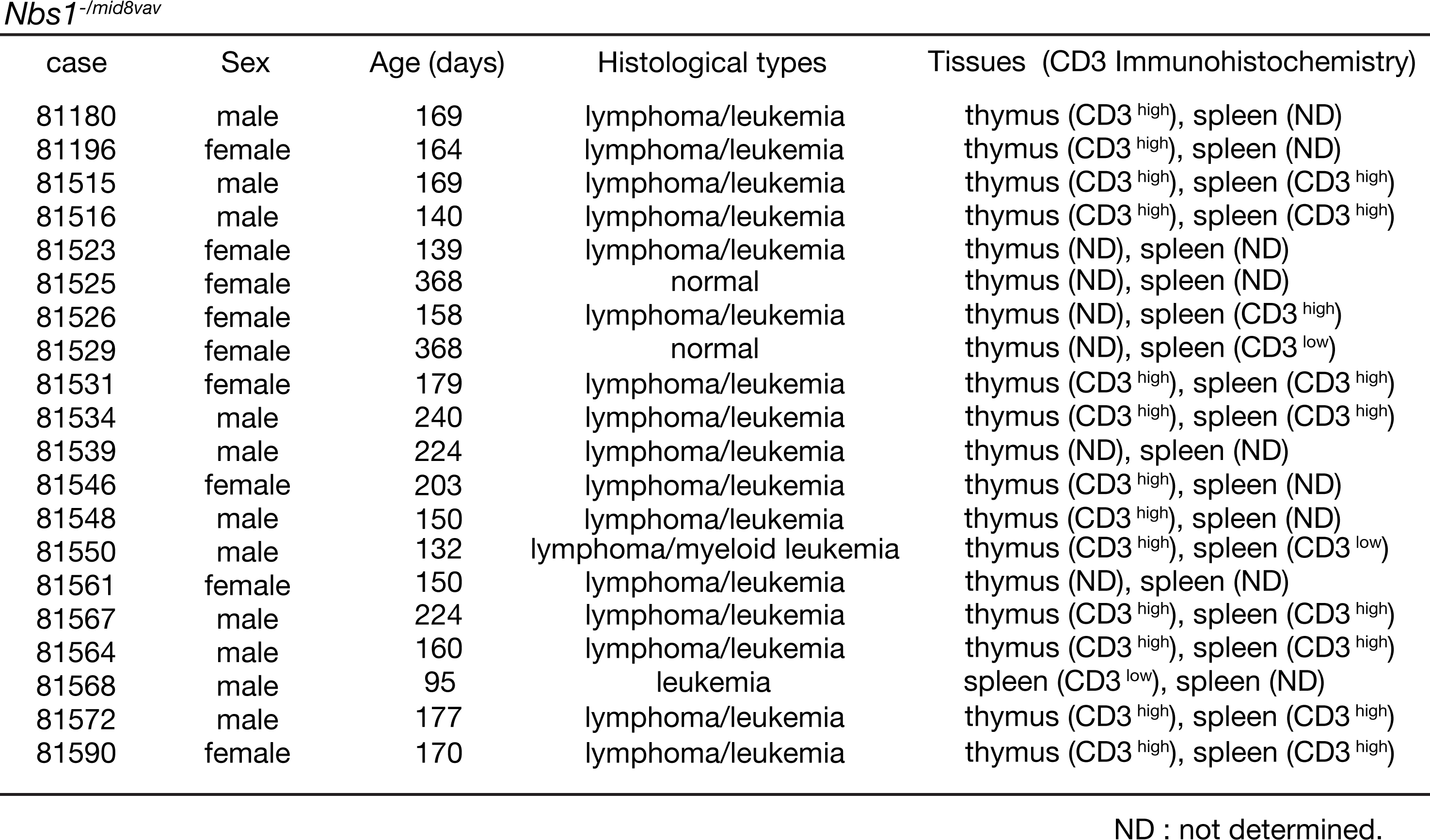
Pathology of *Nbs1*^*-/mid8vav*^mice.

**Figure 3.**
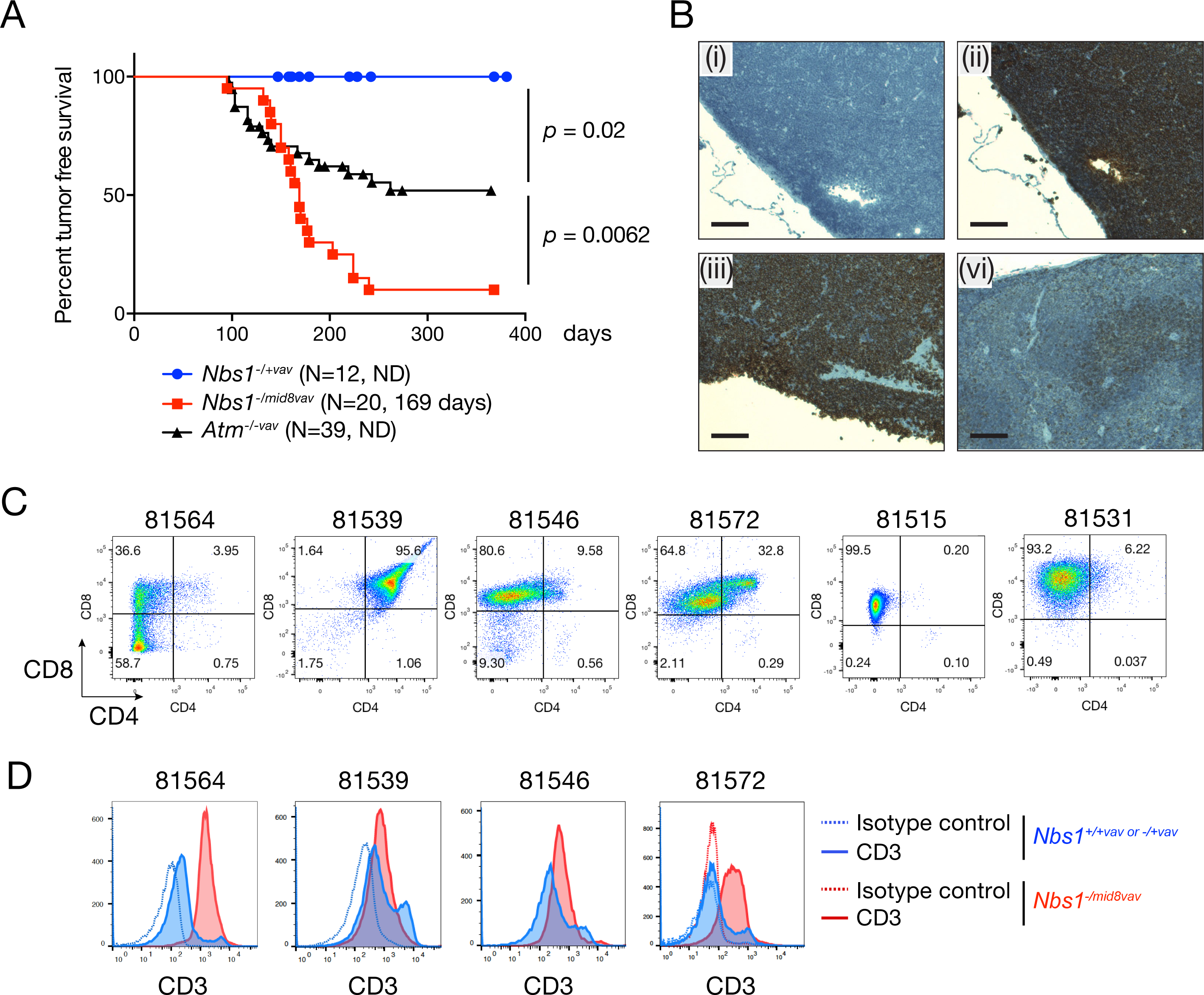
*Nbs1*^*-/mid8vav*^ mice develops spontaneous T cell lymphoma. (A) Mouse tumor free survival. Each data point represents the percent survival of mice with *Nbs1*^*-/+vav*^ and *Nbs1*^*-/mid8vav*^ at a given age. N denotes total number of mice for each genotype and the average age in death in days is shown. ND means not determined. Note that the *Atm*^*-/-vav*^ survival curve was previously established in our colony and reported (Balestrini et al., 2016). (B) CD3 immunohistochemistry of thymus (i and ii) and spleen (iii and vi); control IgG (i), CD3 antibody (ii, iii, and vi), thymus (i and ii) and spleen (iii) from tumor bearing *Nbs1*^*-/mid8vav*^ mouse, and spleen (vi) from littermate *Nbs1*^*-/+vav*^mouse. Scale bar = 100 μm. (C) CD4 and CD8 flow cytometry of thymic tumor cells from *Nbs1*^*-/mid8vav*^ mice. (D) CD3 flow cytometry of thymic tumor cells from *Nbs1*^*-/+vav or +/+vav*^mice and *Nbs1*^*-/mid8vav*^ mice. The numbers above the scatter plots or histograms are the mouse identification number (C and D).

Whereas *Atm*^*-/-vav*^ tumors exhibit a relatively immature phenotype (CD3^low^ CD4^+^ CD8^+^) (Zha et al., 2010), *Nbs1*^*-/mid8vav*^ tumors were predominantly single positive (CD8^+^) in half of the cases examined CD3^high^ (Figure 3C and D). Flow cytometric analysis of thymocyte populations at six weeks of age reveal the CD8^+^ population is elevated in *Nbs1*^*-/mid8vav*^ thymus compared to *Nbs1*^*-/+*^ or *Atm*^*-/-*^ thymus (Figure S1) prior to the emergence of malignant cells.

### Genomic analysis of *Nbs1*^*mid8*^ T cell lymphoma

To gain insight regarding the underlying genetic changes that suppressed the lethality of the *Nbs1*^*mid8*^ allele, we carried out genomic analyses of *Nbs1*^*-/mid8vav*^ tumors. Analysis of copy number variation (CNV) indicated the presence of multiple broad DNA copy number gains and losses, with recurrent CNV on chromosomes 9 and 15 (Figure 4A). A focal amplification of 9qA2 was present in all tumors albeit with variable amplitudes and breakpoints (Figure S2). This region includes the genes encoding *MRE11* and *CHK1*. Chromosome 9 amplification appears to be unique to *Nbs1*^*-/mid8vav*^ tumors as amplification of this region has not been observed in thymic lymphomas from *p53* or *ATM* deficient mice (Dudgeon et al., 2014; Genik et al., 2014; Zha et al., 2010). Thus, the evolution of *Nbs1*^*-/mid8vav*^ malignancy is distinct.

**Figure 4.**
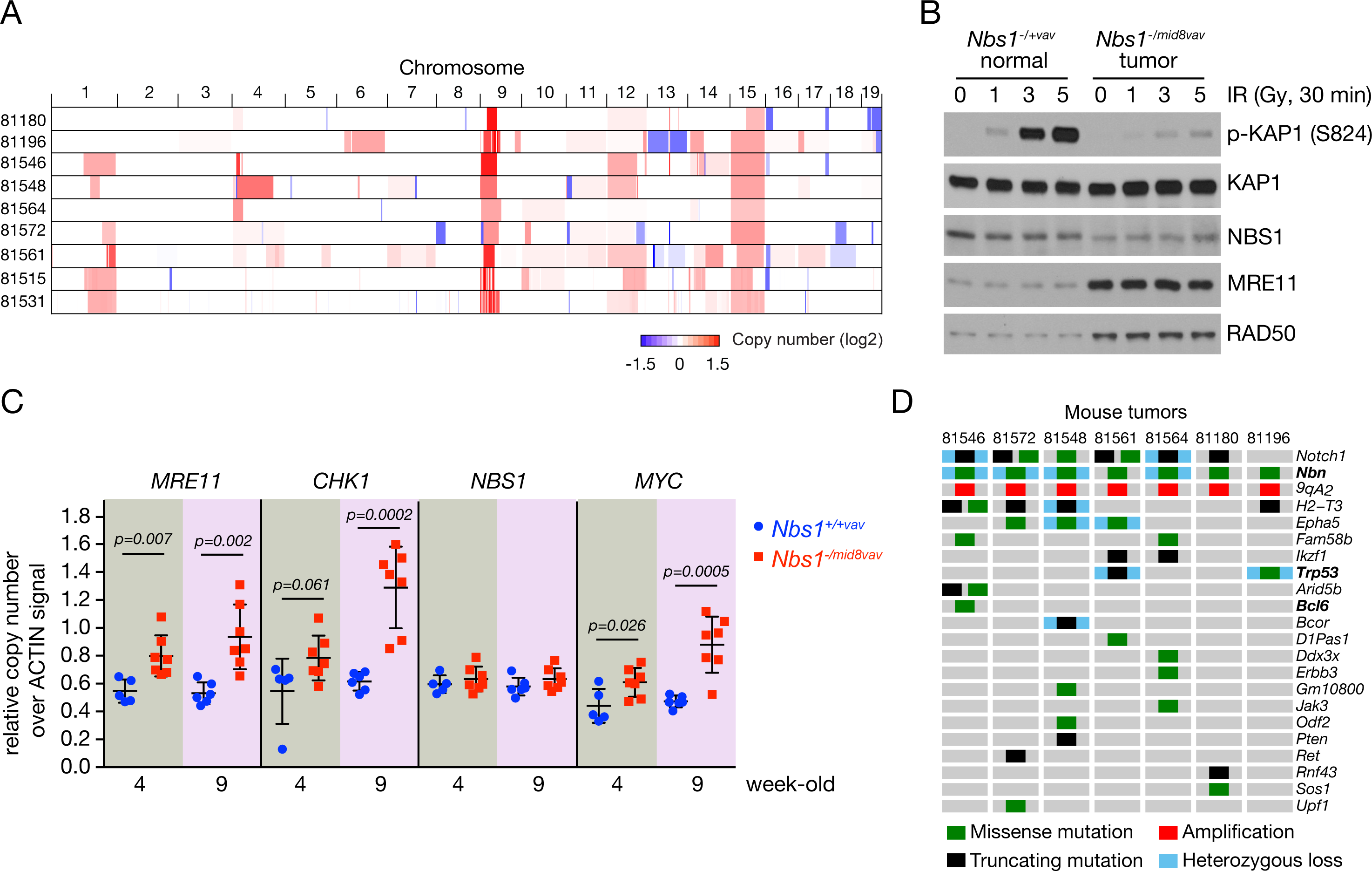
Genomic analysis of *Nbs1*^*-/mid8vav*^ T cell lymphoma. (A) Heapmap of copy number variation (CNV) of 9 thymic tumors lymphomas from *Nbs1*^*-/mid8vav*^ mice. Analysis was performed using whole genome sequencing or deep targeted sequencing of 578 key cancer-associated genes. (B) Western blot for ATM signaling and the levels of Mre11 complex in *Nbs1*^*-/mid8vav*^ thymic tumors. IR-induced phosphorylation of KAP1 (S824) was assessed upon different dose of IR as indicated. The levels of MRE11, NBS1 and RAD50 were assessed. Note that MRE11 and RAD50 levels are increased. (C) CNV of *MRE11, CHK1, NBS1*, and *MYC* genes from thymocytes of wild type and *Nbs1*^*-/mid8vav*^ mice at the age of 4 and 9 weeks. Relative copy numbers for genes over *ACTIN* signals of each biological sample were shown, *n=*5 of *Nbs1*^*+/+vav*^ (4 week-old), *n=*7 of *Nbs1*^*-/mid8vav*^ (4 week-old), *n=*6 of *Nbs1*^*+/+vav*^ (9 week-old), *n=*7 of *Nbs1*^*-/mid8vav*^ (9 week-old), mean ± SD, unpaired t test. (D) Oncoprint of genomic alterations of *Nbs1*^*-/mid8vav*^ T cell lymphoma. Note that *Nbn* mutation is *Nbs1*^*mid8*^ allele (N682_K685del). Complete mutation list is presented in supplemental Table 1. The mouse identification numbers are indicated (A and D).

Amplification was correlated with increased protein levels of Mre11 in *Nbs1*^*-/mid8*^ thymic tumors. Although there was no CNV for the Rad50 gene, Rad50 protein levels were also increased (Figure 4B). We propose that *Nbs1*^*-/mid8vav*^ cells select for amplification and overexpression of Mre11 to compensate for the meta-stable interaction with the Nbs1^mid8^ gene product and thereby permit cell survival. The increased levels of Mre11 in *Nbs1*^*-/mid8vav*^ tumors did not fully suppress the *Nbs1*^*mid8*^ phenotype, as IR-induced Kap1 S824 phosphorylation, an index of ATM activation was not restored (Figure 4B).

In this context, we sought to define when these two gene amplifications occurred in the transition to malignancy. We measured the copy numbers of genes that are present in chromosomes 9 or 15 at earlier times (4 and 9 week-old) prior to the age of tumor onset. Increased copy number of the *MRE11* and *CHK1* genes on chromosome 9 was detected in *Nbs1*^*-/mid8*^ thymocytes as early as four weeks of age (1.46-fold for *MRE11*; 1.48-fold for *CHK1*) while the *NBS1* copy number was not altered (Figure 4C). *MYC* amplification on chromosome 15 was also detected in thymocytes of four week-old *Nbs1*^*-/mid8*^ mice (1.38-fold), suggesting that amplifications on both chromosomes 9 and 15 occurred at an early stage in the leukemogenic process, presumably reflecting selection pressure for cell survival.

The finding of *CHK1* amplification in *Nbs1*^*-/mid8vav*^ T cell leukemias was reminiscent of the fact that human T-ALL commonly exhibits *CHK1* overexpression, and are exquisitely sensitive to Chk1 inhibition (Sarmento et al., 2015). Because the *MRE11* and *CHK1* loci are linked on chromosome 9, it is unclear whether their respective copy number increases are independent events or occurred simultaneously.

Deep targeted sequencing of 578 key cancer-associated genes to identify potential drivers of T-ALL in *Nbs1*^*-/mid8vav*^ mice revealed that the mutational features of *Nbs1*^*-/mid8vav*^ tumors were consistent with those of T-cell acute lymphoblastic leukemia (T-ALL) in humans and are similar, but not identical to lymphomas arising in *Atm*^*-/-*^ mice. *Nbs1*^*-/mid8vav*^ leukemias uniformly contained activating mutations of *Notch1* which are mainly restricted to the PEST domain of the NOTCH1 protein (Table S1 and Figure S3) as seen in human T-ALL (Weng et al., 2004). Amplification or activating mutations in *Notch1* have been also been noted in thymic lymphomas arising in ATM deficient mice (Zha et al., 2010). Those lymphomas also contain trisomy 15 and *Nbs1*^*-/mid8*vav^ tumors similarly appear to have whole chromosome 15 duplications (Figure 4A) (Zha et al., 2010).

Beyond *Notch1*, mutations of other potential driver genes were found at various frequencies in *Nbs1*^*-/mid8vav*^ tumors (Figure 4D and Table S1). *Trp53* mutations within the DNA binding domain were found with a high frequency, and were frequently biallelic due to LOH (Figure 4D and Figure S3). The two mutations observed (S238P corresponding to human S241 and truncation of oligomerization domain) affect DNA binding and p53-mediated cell growth suppression functions (Fischer et al., 2016; Maki, 1999; Monti et al., 2002; Rajagopalan et al., 2010), suggesting that p53 deficiency may be a driver of *Nbs1*^*mid8VAV*^ T-ALL. Other candidate driver mutations found in *Nbs1*^*-/mid8vav*^ tumors including *Arid5b, Bcl6, Bcor*, and *Ikzf1* (Figure 4D and Table S1).

### Overexpression of *MRE11* and *CHK1* rescues the lethality of *Nbs1*^*mid8*^ allele in MEFs

To test the interpretation that Mre11 amplification and overexpression are compensatory changes that can suppress *Nbs1*^*mid8*^ lethality, we overexpressed the murine Mre11 cDNA in *Nbs1*^*flox/mid8*^ creERT2 MEFs (Kim et al., 2017). In these cells, *cre* expression is activated by tamoxifen (4-OHT) to effect inactivation of the (wild type) *Nbs1*^*flox*^ allele to create *Nbs1*^*-/mid8*^ cells. *MRE11*-expressing *Nbs1*^*flox/mid8*^ cells were treated with 4-OHT for 48 hrs, and plated. We were unable to establish viable colonies of *MRE11*-expressing *Nbs1*^*-/mid8*^.

As noted above, the Mre11 complex plays an integral role in DNA replication (Stracker and Petrini, 2011; Syed and Tainer, 2018). Consistent with this interpretation, cells in which the Mre11 complex is impaired experience signs of DNA replication stress (Bruhn et al., 2014). On that basis, we tested the hypothesis that the co-amplification of the *CHK1* locus uniformly observed (Figure 4A and C) is an obligate event for viability of *Nbs1*^*-/mid8vav*^. Therefore, *CHK1* and *MRE11* were co-overexpressed to determine whether this would rescue *Nbs1*^*-/mid8*^ lethality. After deletion of the conditional *Nbs1* allele, cell clones which were *Nbs1*^*-/mid8*^ but overexpressed *CHK1* and *MRE11* emerged (hereafter referred to as *Nbs1*^*-/mid8*^ CM). Three independent *Nbs1*^*-/mid8*^ CM clones (#2, 7,and 8) were verified by PCR genotyping (data not shown) and RT-PCR/sequencing confirmed that the sole source of RNA for Nbs1 protein is *Nbs1*^*mid8*^ mutant allele (Figure S4).

The *Nbs1*^*-/mid8*^ CM clones exhibited slower growth than the parental *Nbs1*^*F/mid8*^ CM cells (Figure 5A). Mre11 and Chk1 levels were increased in three *Nbs1*^*-/mid8*^ CM clones (Figure 5B). We also found that in clone #2, Nbs1 levels were also increased (Figure 5B) due to 7.7-fold amplification of the *Nbs1*^*mid8*^ locus in that clone (Figure 5C), likely in response to selective pressure imposed by the impaired interaction of Mre11 and Nbs1^mid8^.

**Figure 5.**
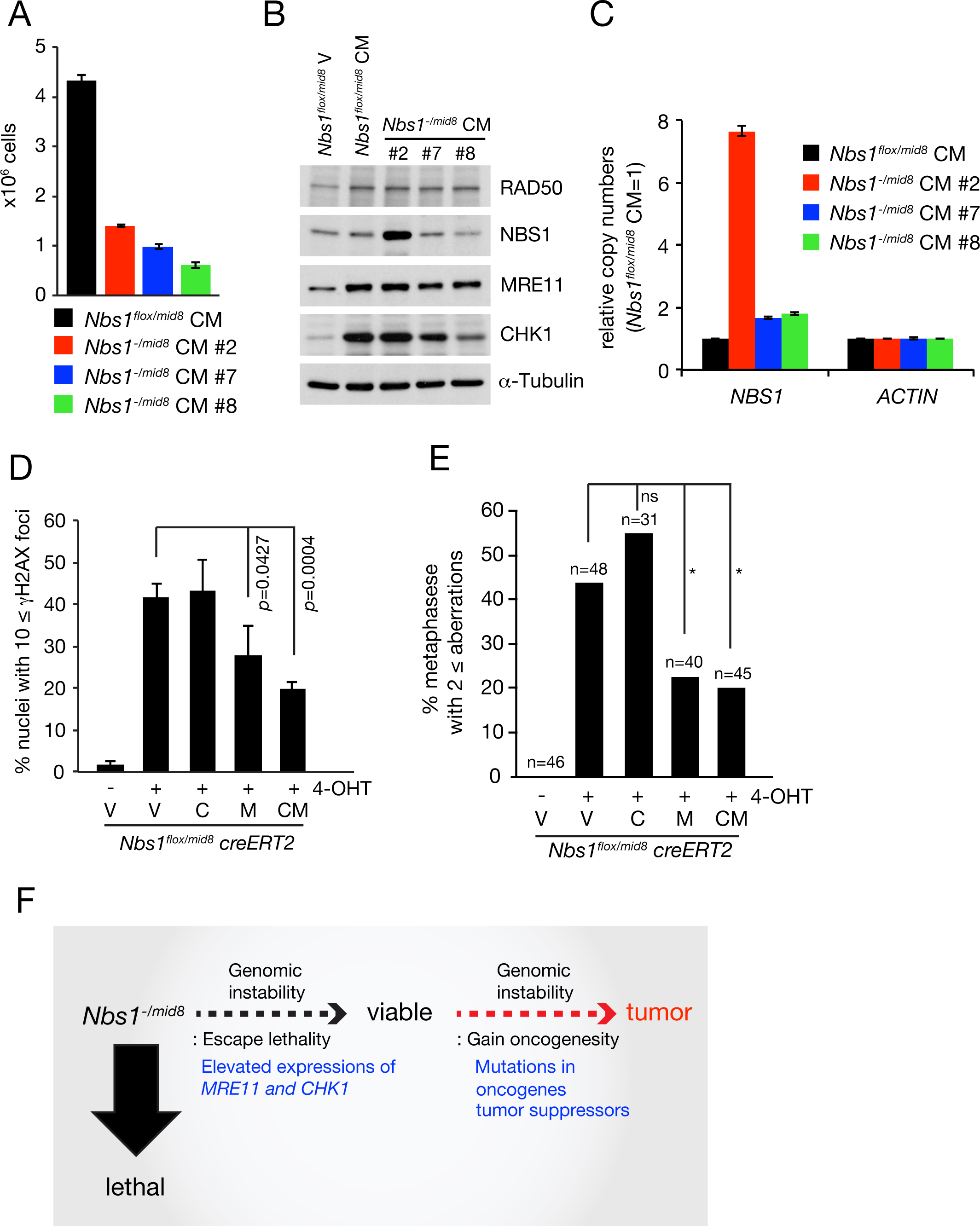
Increased expression of *MRE11* and *CHK1* genes rescues *Nbs1*^*-/mid8*^ MEFs from lethality. (A) Growth of three different stable clones of *Nbs1*^*-/mid8*^ MEFs expressing both exogenous *MRE11* and *CHK1*. *Nbs1*^*flox/mid8*^ CM is a parental cell. Cells were plated (2 x 10^4^) and counted at day 4, mean± SD, triplicates. Hereafter V, C, and M denote empty *<u>V</u>ECTOR, <u>C</u>HK1*, and *<u>M</u>RE11*, respectively. (B) Western blots show the levels of MRE11, RAD50, NBS1, and CHK1 in three stable *Nbs1*^*-/mid8*^ CM clones. *Nbs1*^*flox/mid8*^ CM is a parental cell. (C) *NBS1* gene amplification occurs in clone #2 of *Nbs1*^*-/mid8*^ CM. The values are relative copy numbers of *NBS1* gene compared to that of parental *Nbs1*^*flox/mid8*^ CM cells. (D and E) Genomic instability in *Nbs1*^*-/mid8*^ MEFs was alleviated by ectopic expression of *MRE11*. (D) Percentage nuclei with 10 ≤ γH2AX foci and (E) metaphases with 2 ≤ aberrations were measured in *Nbs1*^*flox/mid8*^ CreERT2 MEFs expressing either *MRE11* and Chk1 alone or in combination, 5 days post 4-OHT treatments. *p* value was determined by unpaired t test, mean ± SD, total more than 800 nuclei counted from 3 independent experiments for γH2AX foci (D) and Fisher’s exact test, \**p*<0.05, ns: not significant (E). (F) Summary of T cell lymphomagenesis in *Nbs1*^*-/mid8vav*^ mice. Gross genomic instability of *Nbs1*^*-/mid8*^ T cells causes impairment in cell proliferation and developmental defects. Few cells that spontaneously gain *MRE11* and *CHK1* amplification escape from the cellular lethality. By acquiring additional tumor-prone mutations, these cells can transform into thymic lymphoma.

To assess whether *MRE11* and *CHK1* overexpression alleviate DNA damage occurring in *Nbs1*^*-/mid8*^ cells, we compared γH2AX foci and metaphase spreads in cells overexpressing either *MRE11* and *CHK1* alone or in combination. Five days after deletion of the conditional *Nbs1*^*flox*^ allele, the ensuing *Nbs1*^*-/mid8*^ cells exhibited multiple indices of DNA damage such as γH2AX foci and chromosomal aberrations (Figure 5D and E). Overexpression of *MRE11* and *CHK1* in combination mitigated those outcomes, reducing γH2AX; 41.8% *vs.* 19.7%, and metaphase aberrations; 43.8% *vs.* 20% (Figure 5D and E). Whereas *CHK1* overexpression alone had no effect, *MRE11* overexpression alone was sufficient to reduce DNA damage in *Nbs1*^*-/mid8*^ cells (Figure 5D and E) even though *MRE11*-expressing *Nbs1*^*-/mid8*^ clones were not ultimately viable. This indicates that although overexpression of *MRE11* was sufficient to alleviate some of genomic instability of *Nbs1*^*mid8*^, coincident *CHK1* overexpression was required for cell viability.

## Discussion

In previous work, we established evidence that the essential function of Nbs1 was to stabilize and promote proper assembly and function of the Mre11 complex using biochemical approaches and atomic force microscopy. In *Nbs1*^*-/mid8*^ cells, Mre11 complex components are present at essentially normal levels, but Nbs1^mid8^ interaction with Mre11 is compromised, blocking viability. Four days after inactivation of the wild type allele in *Nbs1*^*flox/mid8*^ MEFs (prior to death), ATM activation was abolished and severe genomic instability was evident (Kim et al., 2017). In this study, we used the context of hematopoietic development to assess whether Nbs1^mid8^ retained residual function.

Whereas Nbs1 deficiency in the bone marrow led to peri-natal lethality due to anemia, *Nbs1*^*-/mid8vav*^ mice were viable with a median lifespan of 169 days. *Nbs1*^*-/mid8vav*^ succumbed to a highly aggressive T cell malignancy that bore some hallmarks of T-ALL in humans. Genomic analysis of *Nbs1*^*-/mid8vav*^ cancers revealed copy number gains and overexpression of *MRE11* and *CHK1* in 100% of tumors analyzed. We were able to establish clonal *Nbs1*^*-/mid8*^ MEF cell lines only upon co-overexpression of *MRE11* and *CHK1*, demonstrating that increased levels of both was required to suppress *Nbs1*^*mid8*^ lethality.

### Mre11 Complex Assembly and Tumor Suppression

Each member of the Mre11 complex is essential and none appear to function outside of the complex. Nevertheless, Nbs1 plays a distinct role in Mre11 complex stability. For example, mutations that reduce Nbs1 levels do not grossly alter the levels of Mre11 and Rad50 (Carney et al., 1998; Williams et al., 2002), whereas the stabilities of Mre11 and Rad50 are interdependent, so that mutations destabilizing either reduce the level of the other (Stewart et al., 1999; Theunissen et al., 2003). Hence, Mre11 and Rad50, which are present in all clades of life constitute the core of the Mre11 complex, while Nbs1, seen only in Eukarya has evolved to promote proper assembly and activity of the complex in eukaryotes.

We propose that this function of Nbs1 underlies the selection for *MRE11* amplification and overexpression in *Nbs1*^*-/mid8vav*^ tumors. Based on principles of ligand binding equilibria, the weakened interaction between Nbs1^mid8^ and Mre11 would be partially mitigated if the levels of either Mre11 or Nbs1^mid8^ were elevated. This could increase the steady state level of productively assembled complexes to a threshold that would permit viability. Supporting this concept, in *Nbs1*^*-/mid8*^ CM clone 2, the *Nbs1*^*mid8*^ locus was also amplified and overexpressed (Figure 5B and C). This suppression is not complete, as the defect in Kap1 S824 phosphorylation upon IR treatment of *Nbs1*^*-/mid8vav*^ tumors was not rescued (Figure 4B). We interpret this to mean that the levels of putatively assembled complexes were insufficient to allow for IR-induced ATM activation.

Several mouse models of Mre11 complex hypomorphism have been described (Roset et al., 2014; Stracker and Petrini, 2011). No predisposition to malignancy has been observed in these mutants, although in some cases the latency of tumorigenesis associated with p53 or Chk2 mutations is reduced; presumably this reflects the combined effect of genome instability and checkpoint defects associated with those mutations.

The strong cancer predisposition seen in *Nbs1*^*-/mid8vav*^ is distinct from other Mre11 complex single mutants, and phenocopies that of *Nbs1*^*ΔB/ΔB*^ *Atm*^*-/-vav*^ mice. Two features of these mice likely underlie the similar outcomes (Balestrini et al., 2016). First, ATM activity is virtually absent in the former and completely absent in the latter, which means that in both contexts, DNA damage dependent cell cycle checkpoints and apoptotic induction are compromised. Second, both exhibit high degrees of genomic instability, which likely increases the probability of chromosome rearrangement and/or mutations that enhance progression to the malignant phenotype.

### The Requirement for *CHK1* Overexpression

In *Nbs1*^*-/mid8*^ MEFs, overexpression of *MRE11* alone is not sufficient to restore viability. Co-overexpression of *CHK1* was required to obtain stable clones (Figure 5B). This finding strongly suggests that amplification and overexpression of the *MRE11* and *CHK1* loci in *Nbs1*^*-/mid8vav*^ T-ALL cells reflects the same requirement. Significantly increased copy numbers of both loci were observed as early as four weeks of age, supporting the interpretation that the overexpression of both genes is required for viability irrespective of the malignant phenotype.

Whereas *MRE11* overexpression in *Nbs1*^*-/mid8*^ MEFs mitigated the gross chromosome instability observed after *cre* meditated deletion of the *Nbs1*^*flox*^ allele (Figure 5E), the requirement for *CHK1* overexpression suggests an additional stress is imposed by the Nbs1^mid8^-containing Mre11 complex. It is likely that with the ATM-Mre11 complex axis of the DDR crippled by the *Nbs1*^*mid8*^ allele, selection for *CHK1* overexpression is imposed by the genomic instability associated with severe Mre11 complex hypomorphism. In this scenario, Chk1 function may mitigate the lethal effects of DNA damage in parallel with its normal role in the response to DNA replication stress.

DNA replication stress, broadly defined as a state in which progression of the replisome is impaired by DNA lesions or insufficiency of the nucleotide pool is an important source of DNA damage in proliferating cells. DNA replication stress also appears to be intrinsic to the premalignant and malignant phenotypes (Gaillard et al., 2015; Macheret and Halazonetis, 2015). The viability of cells experiencing DNA replication stress is largely dependent on the ATR-Chk1 axis of the DNA damage response (Schoppy et al., 2012; Zeman and Cimprich, 2014), and so the development of Chk1 and ATR inhibitors has emerged as a priority in cancer therapeutics (Pilie et al., 2018).

Given its requirement for preserving genomic integrity during S phase (Adelman et al., 2009), it is also conceivable that the selection for *CHK1* overexpression reflects that Mre11 complex depletion causes DNA replication stress in a manner analogous to depletion of MCM proteins and other replisome components (Hills and Diffley, 2014), a state that acutely requires Chk1 activity to maintain viability (Gaillard et al., 2015; Schoppy et al., 2012).

Cytologic analyses of acute Nbs1 depletion in MEFs offered the suggestion that the Mre11 complex’s role in resolving DNA replication intermediates underlies its essentiality (Bruhn et al., 2014). Perhaps consistent with that view, we have shown that Mre11 complex foci arise during unperturbed S phase. Those foci do not colocalize with DSB markers such as γH2AX or BRCA1, but colocalize with PCNA throughout S phase (Maser et al., 2001; Mirzoeva and Petrini, 2003). Hence, they do not appear to be associated with DSBs and could potentially represent DNA replication intermediates that precede fork collapse and DSB formation.

An additional possibility is that the Mre11 complex is situated at the fork to degrade secondary structures on the lagging strand and thereby drive sister chromatid recombination as has been suggested in bacteria and *S. cerevisiae* (Connelly and F., 1996; Connelly et al., 1998; Lobachev et al., 2002). Whatever the source of the initiating DSBs, the Mre11 complex is acutely required for sister chromatid recombination (Stracker and Petrini, 2011), a function that could also account for Chk1 activation upon depletion as well as the complex’s association with the fork under normal conditions.

### *Nbs1*^*-/mid8vav*^ and T-ALL in Humans

Genomic analysis of *Nbs1*^*mid8*^ leukemias revealed a similar patter of mutations to that found in human T-ALL (Sarmento et al., 2015; Weng et al., 2004). The *Notch1* mutations were found in 6 tumors out of 7 samples analyzed and the mutations were restricted to the PEST domain, as seen in human T-ALL and also T-ALL arising in *Atm*^*-/-vav*^ mice (Weng et al., 2004; Zha et al., 2010). This suggests that *Notch1* mutations could be the main driver for T-ALL tumors. Other mutations found in *Nbs1*^*-/mid8vav*^ leukemias are also known human tumor mutations. For example, *Bcl6, Bcor, and Ikzf1* are found mutated in human T-ALL and B-ALL (Kastner and Chan, 2011; Mullighan et al., 2008). *Bcl6* is a regulator for germinal center reaction and found deregulated in ~40 % diffuse large B-cell lymphomas (DLBCL) by translocation (Butler et al., 2002; Lo Coco et al., 1994; Ye et al., 1993). Loss of function mutations in *Bcor* has been noted in hematopoietic malignancies (Damm et al., 2013; Grossmann et al., 2011). In this regard, *Nbs1*^*-/mid8vav*^ may provide an *in vivo* model for examining the contributions of the various driver mutations in T-ALL and other malignancies in the context of Mre11 complex hypomorphism.

The Mre11 complex plays a critical role in preventing oncogene induced carcinogenesis in mammary epithelium (Gupta et al., 2013). It is conceivable that Mre11 complex hypomorphism in *Nbs1*^*-/mid8vav*^ cells similarly creates a permissive state for oncogene driven proliferation and malignancy. The data presented here reveal that Nbs1 meets the definition of a tumor suppressor in hematopoietic cells via its role in ATM activation and in DNA repair. Therefore, the tumor suppressive functions defined in the epithelium appear to be recapitulated in the hematopoietic compartment. Further examination of genetic interactions between Mre11 complex hypomorphism and oncogenic driver mutations in hematologic malignancy will shed light on the mechanisms underlying the tumor suppressive role of the Mre11 complex in this context.

## Materials and methods

### Mice

The generation of *Nbs1*^*mid8*^, *Nbs1*^*flox/flox*^ *vav*^*cre*^, and *Atm*^*flox/flox*^ *vav*^*cre*^ mice were previously described (Balestrini et al., 2016; Kim et al., 2017). *Nbs1*^*flox/mid8*^ *vav*^*cre*^ mice were generated by crossing with *Nbs1*^*flox*^ and *vav*^*cre*^ mice. For tumor watch, the cohorts were maintained according to the protocol approved by the Institutional Animal Care and Use Committee of Memorial Sloan Kettering Cancer Center. Tumor samples from the sacrificed mice were fixed with 4% (v/v) formaldehyde overnight at 4°C and stored at 4°C in 70% ethanol. CD3 antibody (ab16669, Abcam) was used for immunohistochemistry. Eight micrometer sections of paraffin-embedded samples were stained with Hematoxylin and Eosin (H&E) for pathological analysis. Histopathology was performed by Histoserv, Inc. (Maryland, USA).

### Cell lines

Mouse embryo fibroblasts (MEFs) were cultured in DMEM/10% cosmic calf serum. Inducible *Nbs1*^*flox/mid8*^ MEFs were generated by stable expression of MSCV CreERT2 puro (a gift from Tyler Jacks; Addgene plasmid #22776). Stable *CHK1* or/and *MRE11* expressing *Nbs1*^*flox/mid8*^ CreERT2 MEFs were made by retroviral expression of *CHK1* or *MRE11* in pMIG-W-IRES-GFP plasmid (a gift from Luk Parjis; Addgene plasmid #12282). Then, inducible *Nbs1* allele was removed by 4-OHT treatment. After deletion of inducible *Nbs1* allele, individual surviving clones were tested by PCR genotyping and Sanger sequencing of amplified region by RT-PCR to verify absence of wild type *Nbs1* allele expression. Three independent *Nbs1*^*-/mid8*^ *CHK1*+*MRE11* MEFs clones were established.

### Peripheral blood analysis

Peripheral blood was collected from the tail vein. A compete peripheral blood count was measured using a Hemavet 950 (Drew Scientific).

### Flow cytometry analyses

Thymus and femurs from 6-8 week old mice were harvested and subjected to red blood cell lysis (00-4333-57, Invitrogen). Thymocytes and bone marrow cells were counted by Beckman Coulter Z1 counter. Antibodies used for flow cytometry are CD8 (553030, BD Bioscience), CD4 (557308, BD Bioscience), CD3 (100214, Biolegend), B220 (562922, BD Bioscience, 103212, Biolegend), CD43 (553270, BD Bioscience), IgM (562032, BD Bioscience), Mac-1 (557397, BD Bioscience), Gr-1 (553126, BD Bioscience), Ter119 (116223, Biolegend), and Isotype ctrl (400120, Biolegend).

### Sequencing and analysis

Tumor samples along with normal samples taken from the tails of unaffected littermates underwent either a custom-capture and deep sequencing assay of 578 established cancer genes called Mouse Integrated Mutation Profiling of Actionable Cancer Targets (M-IMPACT) or low-pass whole-genome sequencing (WGS) for additional DNA copy number analyses. Tumors were sequenced with M-IMPACT to an average 300.9-fold coverage. Analysis for somatic point mutations and small insertions and deletions along with DNA copy number changes in these data was performed a previously described (Pronier et al., 2018). Tumors were sequenced by low-pass WGS to an average of 4.9-fold coverage. DNA copy number alterations were identified summarizing read coverage in 50kb non-overlapping windows across the genome followed by GC-content normalization using a lowess fit. We then performed circular binary segmentation (alpha of 1e-5 and undo.SD of 5) (Venkatraman and Olshen, 2007). Structural variants were identified using delly v0.6.1 (Rausch et al., 2012).

### Cellular assays

For metaphase spread, cells were treated with 100 ng/ml of KaryoMAX colcemid (Life technologies) for 1 hr and harvested. Cells were hypotonically swelled in 0.075 M KCl for 15 min at 37°C and fixed in ice-cold 3:1(v/v) methanol: acetic acid. Dropped samples on slides were stained with 5% Giemsa (Sigma) and mounted with Permount medium (Fisher Scientific). Analyses were carried out on blinded samples.

For immunofluorescence staining, cells were fixed in 4% (v/v) paraformaldehyde for 15 min at room temperature and permeabilized in 0.5% (v/v) TritonX-100 for 10 min at room temperature before primary antibody incubation. Phospho-H2AX (Ser139) antibody was used (05-636, Millipore) at 1:5000 dilution and secondary antibody (Alexa-594, Life technologies) was used at 1:1000 dilution. Staining PBS buffer contains 1% BSA and 0.1% (v/v) Tween 20.

### Western Blot

Western blots were performed by standard protocol. Total cells extracts were prepared in SDS lysis buffer (60 mM Tris-HCl pH 6.8, 2% SDS) and boiled for 5 min to shear genomic DNA. Twenty microgram of extracts were analyzed for western blot. Antibodies used in this study were MRE11 (custom made), NBS1 (custom made), RAD50 (custom made), p-KAP1 S824 (ab70369, Abcam), total KAP1 (NB500-159, Novus), CHK1 (sc-8408, Santa Cruz), and a-TUBULIN (T9026, Sigma).

### Copy number qPCR

Genomic DNAs were isolated using GeneJET Genomic DNA Purification Kit (Thermo) and qPCRs were performed using SsoAdvanced™ Universal SYBR^®^ Green Supermix (Bio Rad). Specific mouse primer set sequences are *MRE11*; 5’-TCACACTGGATTGGATCTTCCAAGTG-3’, 5’-CTATAGAGTGAGATCCAGAACAGCCA-3’, *NBS1*; 5’-GCCTTCTAGGTGGACATGCT-3’, 5’-TCTGGCACCTGAACTTTGTGT-3’, *Chk1*; 5’-ATCCTTACCTCCAGCCAACATTGCAGT-3’, 5’-ACGCTTACTGAACAAGATGTGTGGGACT-3’, *MYC*; 5-GCTGTTTGAAGGCTGGATTTCCT-3, 5’-CAGATGAAATAGGGCTGTACGGAGTC-3’, *ACTIN*; 5’-CCTCTACATTCATTGTAGGCCACCACA-3’, 5’-CTTCTGAGTTTGGGATGCTGTGAGTG-3’).

## Acknowledgements

We thank Fred Alt for *ATM*^*flox*^ mice, Matthias Stadtfeld and Thomas Graf for *vav*^*cre*^ mice, the MSK Integrated Genomics Operation, and members of the Petrini lab, including Thomas J. Kelly for helpful discussion throughout the course of this study. This work was supported by GM59413 (J.H.J.P), U54 OD020355 (J.H.J.P and B.S.T.), R01 CA207244 (B.S.T.), R01 CA204749 (B.S.T.), and the American Cancer Society (RSG-15-067-01-TBG), Anna Fuller Fund, and the Josie Robertson Foundation (B.S.T.), the Cycle for Survival, and the MSK Cancer Center core grant P30 CA008748.

All authors declare no competing financial interests.

## Author contribution

J. Kim performed experiments; J. Kim, A. Pension, B. Taylor, and J. Petrini analyzed the data and wrote the paper; J. Kim, and J. Petrini designed the study.

---

**Figure S1.**
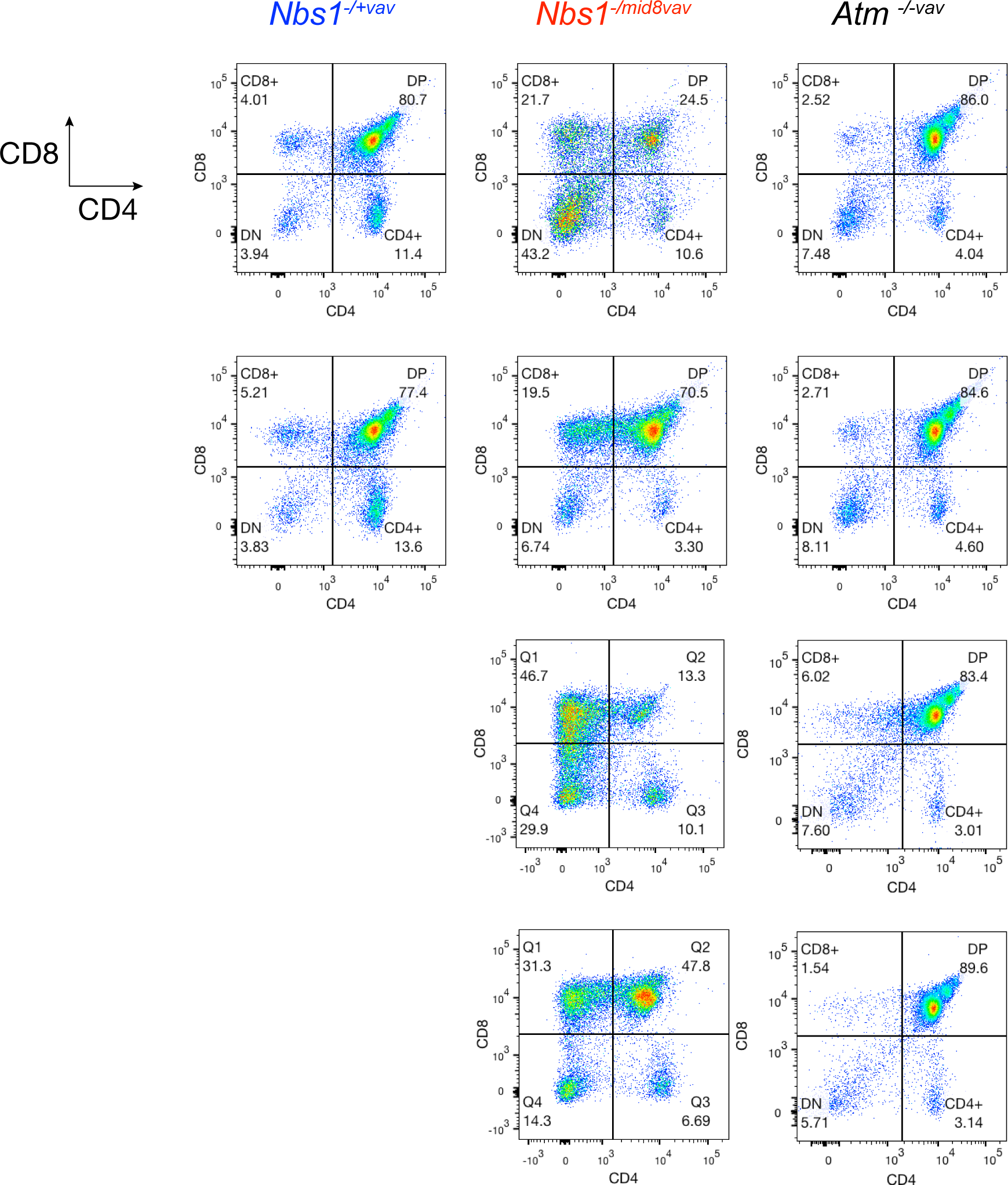
Normal thymocyte analysis in *Nbs1*^*-/mid8vav*^ mice. Thymocytes were isolated from thymus of 6 week-old mice, and analyzed by CD4 and CD8 flow cytometry, n=2 of *Nbs1*^*-/+vav*^, n=4 of *Nbs1*^*-/mid8vav*^, n=4 of *Atm*^*-/-vav*^.

**Figure S2.**
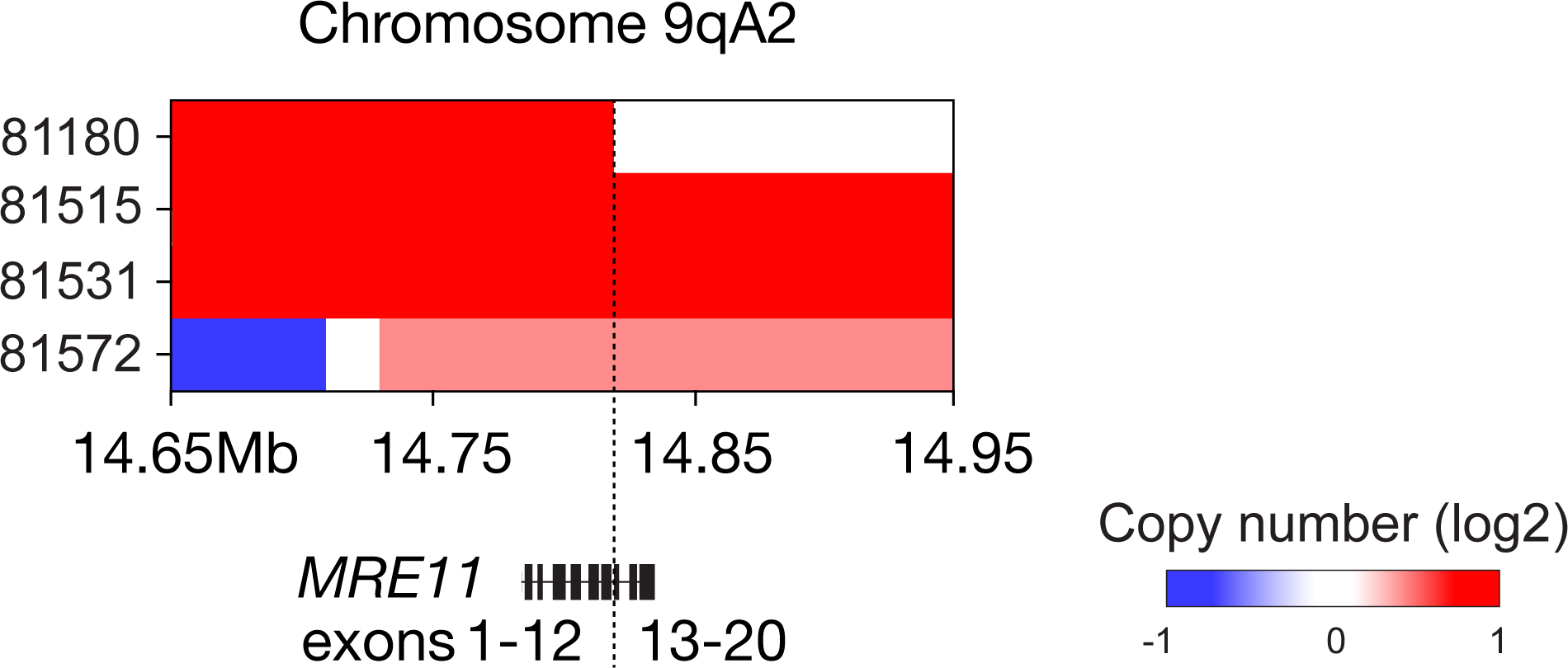
A focal amplification of 9qA2 spanning *MRE11*.

**Figure S3.**
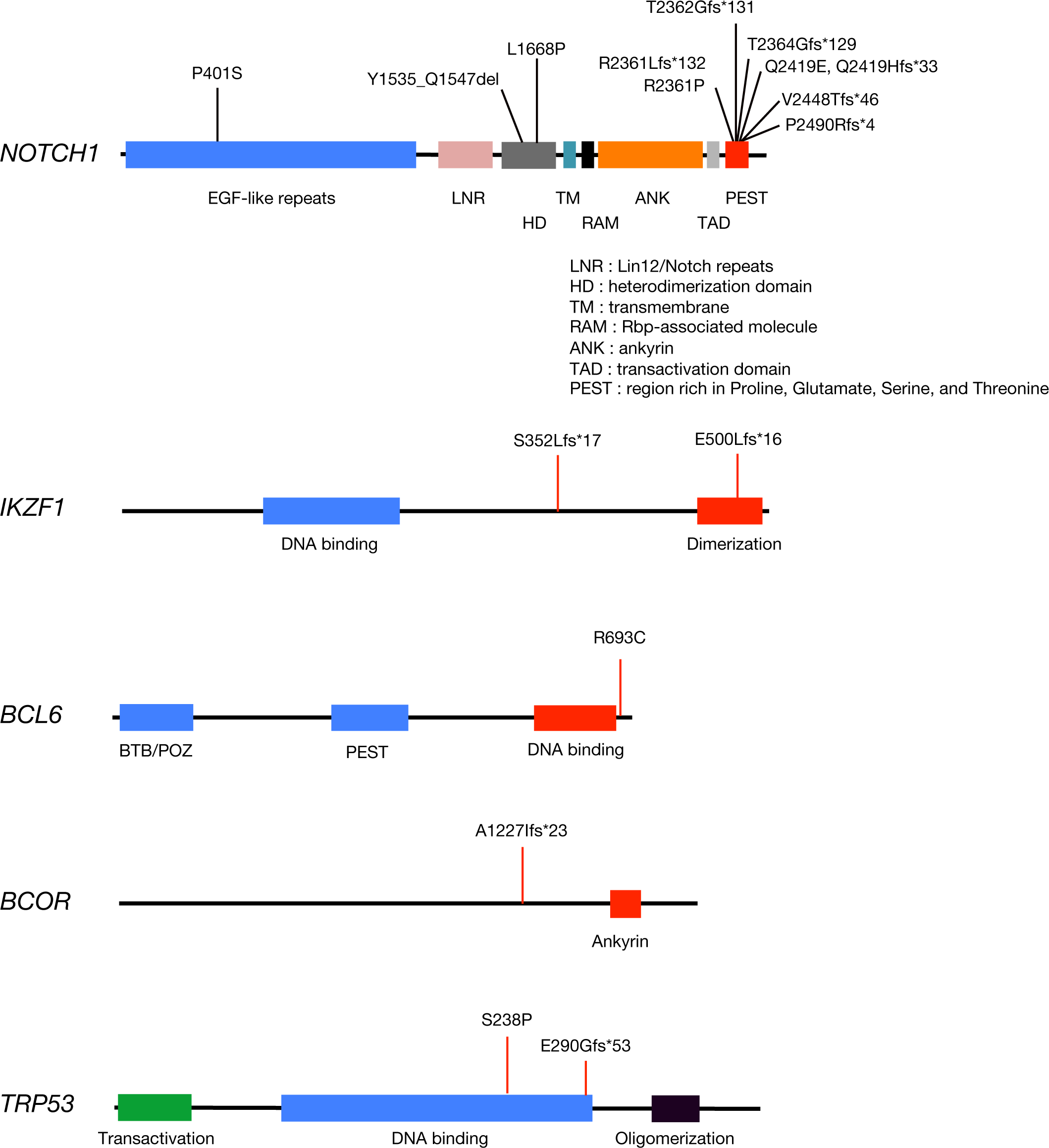
Domain structures of mutation genes found in *Nbs1*^*-/mid8vav*^ tumors.

**Figure S4.**
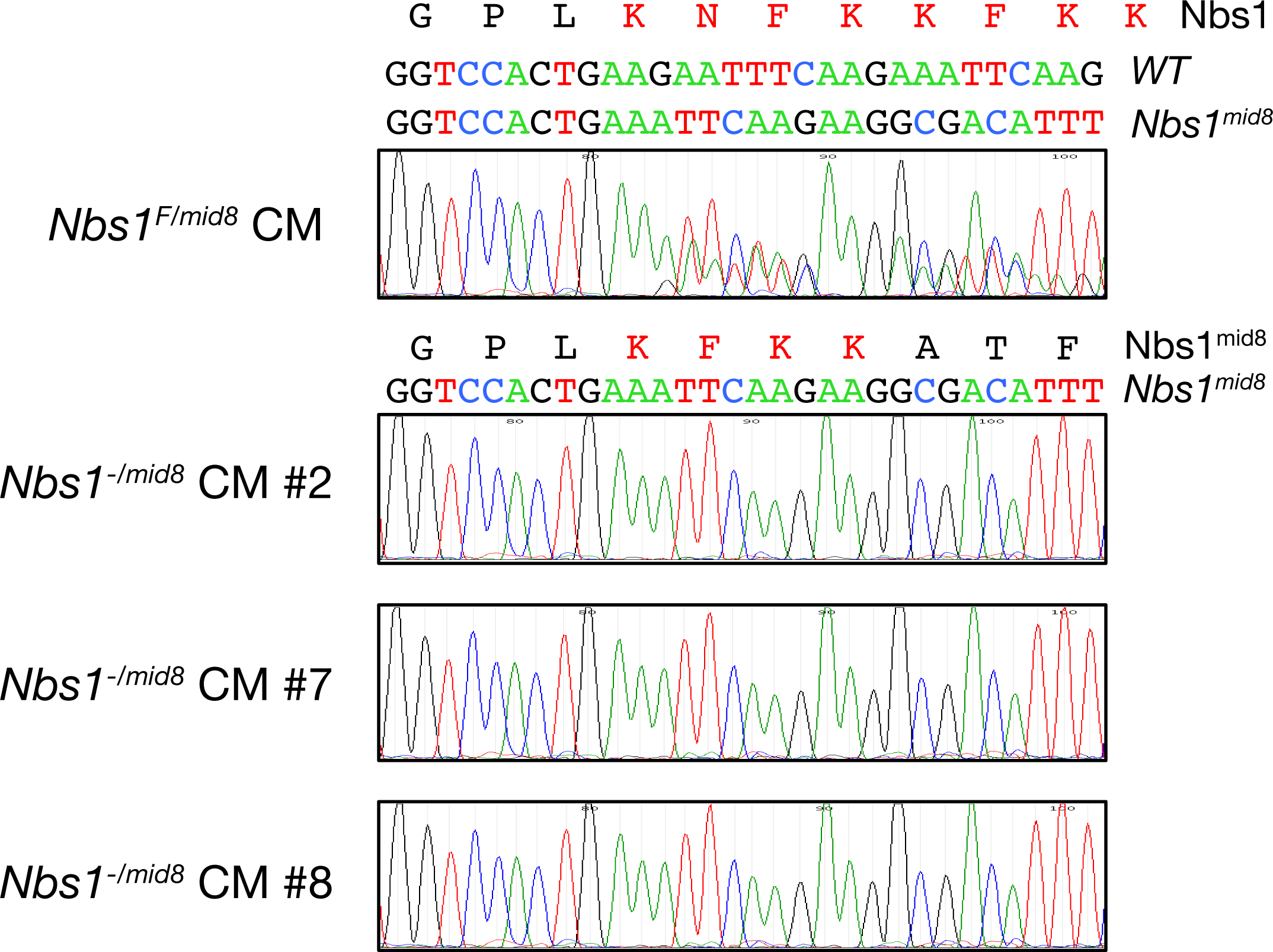
Verification of *Nbs1*^*-/mid8*^ CM clones #2, #7, and #8. The RNA level of *Nbs1*^*mid8*^ allele from three *Nbs1*^*-/mid8*^ CM clones #2, #7, and #8 were confirmed by sequencing the amplified region by RT-PCR.

**Table S1.** Mutations of *Nbs1*^*-/mid8vav*^ tumor samples.

| Hugo_Symbol | Chromosome | Tumor sample | dbSNP_RS | HGVSp_Short | t_depth | t_ref_count | t_alt_count | Variation frequency |
| --- | --- | --- | --- | --- | --- | --- | --- | --- |
| *D1Pas1* | 1 | s_81561 | novel | p.V419M | 127 | 101 | 26 | 0.20 |
| *Notch1* | 2 | s_81572 | novel | p.P2490Rfs*4 | 433 | 370 | 63 | 0.15 |
| *Notch1* | 2 | s_81572 | novel | p.V2448Tfs*46 | 581 | 515 | 66 | 0.11 |
| *Notch1* | 2 | s_81564 | novel | p.Q2419Hfs*33 | 738 | 296 | 442 | 0.60 |
| *Notch1* | 2 | s_81564 | novel | p.Q2419E | 652 | 254 | 398 | 0.61 |
| *Notch1* | 2 | s_81546 | novel | p.L2362Gfs*131 | 509 | 46 | 463 | 0.91 |
| *Notch1* | 2 | s_81561 | novel | p.T2364Gfs*129 | 260 | 145 | 115 | 0.44 |
| *Notch1* | 2 | s_81561 | novel | p.R2361P | 214 | 119 | 95 | 0.44 |
| *Notch1* | 2 | s_81180 | novel | p.R2361Lfs*132 | 670 | 367 | 303 | 0.45 |
| *Notch1* | 2 | s_81548 | novel | p.L1668P | 758 | 717 | 41 | 0.05 |
| *Notch1* | 2 | s_81572 | novel | p.L1668P | 510 | 488 | 21 | 0.04 |
| *Notch1* | 2 | s_81572 | novel | p.Y1535_Q1547del | 552 | 482 | 70 | 0.13 |
| *Notch1* | 2 | s_81572 | novel | p.P401S | 367 | 208 | 158 | 0.43 |
| *Odf2* | 2 | s_81548 | novel | p.D240E | 72 | 66 | 6 | 0.08 |
| *Gm10800* | 2 | s_81548 | novel | p.Q204L | 630 | 620 | 9 | 0.01 |
| *Nbn* | 4 | s_81180 | novel | p.N682_K685del | 416 | 202 | 214 | 0.51 |
| *Nbn* | 4 | s_81196 | novel | p.N682_K685del | 232 | 121 | 111 | 0.48 |
| *Nbn* | 4 | s_81546 | novel | p.N682_K685del | 617 | 154 | 463 | 0.75 |
| *Nbn* | 4 | s_81548 | novel | p.N682_K685del | 479 | 126 | 353 | 0.74 |
| *Nbn* | 4 | s_81561 | novel | p.N682_K685del | 181 | 113 | 68 | 0.38 |
| *Nbn* | 4 | s_81564 | novel | p.N682_K685del | 578 | 92 | 486 | 0.84 |
| *Nbn* | 4 | s_81572 | novel | p.N682_K685del | 314 | 40 | 274 | 0.87 |
| *Epha5* | 5 | s_81548 | novel | p.E298K | 626 | 0 | 625 | 1.00 |
| *Epha5* | 5 | s_81561 | novel | p.E298K | 621 | 1 | 620 | 1.00 |
| *Epha5* | 5 | s_81572 | novel | p.E298K | 542 | 267 | 274 | 0.51 |
| *Ret* | 6 | s_81572 | novel | p.R898* | 762 | 407 | 354 | 0.46 |
| *Upf1* | 8 | s_81572 | novel | p.R402C | 605 | 563 | 42 | 0.07 |
| *Jak3* | 8 | s_81564 | novel | p.A617T | 253 | 248 | 4 | 0.02 |
| *Arid5b* | 9 | s_81546 | novel | p.S1015Rfs*64 | 827 | 661 | 166 | 0.20 |
| *Arid5b* | 10 | s_81546 | novel | p.Y832C | 819 | 759 | 59 | 0.07 |
| *Arid5b* | 10 | s_81546 | novel | p.D798Y | 767 | 669 | 98 | 0.13 |
| *Arid5b* | 10 | s_81546 | novel | p.A797T | 762 | 659 | 103 | 0.14 |
| *Arid5b* | 10 | s_81546 | novel | p.H784Mfs*27 | 1034 | 783 | 251 | 0.24 |
| *Arid5b* | 10 | s_81546 | novel | p.M666V | 904 | 666 | 238 | 0.26 |
| *Arid5b* | 10 | s_81546 | novel | p.A869S | 1003 | 869 | 109 | 0.11 |
| *ErbB3* | 10 | s_81564 | novel | p.L1163F | 428 | 240 | 188 | 0.44 |
| *Ikzf1* | 11 | s_81564 | novel | p.S352Lfs*17 | 875 | 480 | 395 | 0.45 |
| *Ikzf1* | 11 | s_81561 | novel | p.E500Lfs*16 | 1090 | 623 | 467 | 0.43 |
| *Trp53* | 11 | s_81196 | novel | p.S238P | 896 | 12 | 884 | 0.99 |
| *Trp53* | 11 | s_81561 | novel | p.E290Gfs*53 | 675 | 94 | 581 | 0.86 |
| *Fam58b* | 11 | s_81546 | novel | p.N234S | 467 | 459 | 8 | 0.02 |
| *Fam58b* | 11 | s_81546 | novel | p.A211V | 460 | 447 | 13 | 0.03 |
| *Fam58b* | 11 | s_81564 | novel | p.A211V | 566 | 549 | 16 | 0.03 |
| *Rnf43* | 11 | s_81180 | novel | p.W13* | 537 | 525 | 12 | 0.02 |
| *Bcl6* | 16 | s_81546 | novel | p.R693C | 462 | 271 | 190 | 0.41 |
| *H2-T3* | 17 | s_81546 | novel | p.E290K | 7 | 4 | 3 | 0.43 |
| *H2-T3* | 17 | s_81548 | novel | p.E290K | 5 | 2 | 3 | 0.60 |
| *H2-T3* | 17 | s_81196 | novel | p.X209_splice | 13 | 6 | 7 | 0.54 |
| *H2-T3* | 17 | s_81546 | novel | p.X209_splice | 8 | 4 | 4 | 0.50 |
| *H2-T3* | 17 | s_81548 | novel | p.X209_splice | 12 | 3 | 9 | 0.75 |
| *H2-T3* | 17 | s_81572 | novel | p.X209_splice | 12 | 6 | 6 | 0.50 |
| *Sos1* | 17 | s_81180 | novel | p.A1233V | 1149 | 818 | 330 | 0.29 |
| *Pten* | 19 | s_81548 | novel | p.X70_splice | 81 | 64 | 17 | 0.21 |
| *Pten* | 19 | s_81548 | novel | p.X79_splice | 80 | 65 | 15 | 0.19 |
| *Bcor* | X | s_81548 | novel | p.A1227Lfs*23 | 270 | 11 | 259 | 0.96 |
| *Ddx3x* | X | s_81564 | novel | p.R218G | 145 | 121 | 24 | 0.17 |
| *Ddx3x* | X | s_81564 | novel | p.N509I | 223 | 219 | 4 | 0.02 |

